# Mathematical modeling of metastasis, a feasible way to detect the weakness

**DOI:** 10.1101/2020.08.19.257931

**Authors:** A. Guerra, E. Silva, R. Mansilla, J. M. Nieto-Villar

**Author notes:** **Correspondence to:** Prof./Dr. J.M. Nieto-Villar, Department of Chemical-Physics, A. Alzola Group of Thermodynamics of Complex Systems of M.V. Lomonosov Chair, Faculty of Chemistry, University of Havana, Cuba.

## Abstract

**Aim:** Cancer is one of the main causes of death worldwide. 90% of deaths caused by this disease occur due to metastasis. Two models are proposed that rescue fundamental aspects of metastasis, such as EMT (epithelial-mesenchymal transition), extravasation and colonization.

**Methods:** To evaluate the complexity, the Lyapunov exponents, the eigenvalues of the Jacobian matrix (stability analysis) and the Kaplan York dimension were calculated.

**Results:** It was evidenced that the weakness of the metastasis lies in these stages, which indicates that they constitute potential targets in the search for an effective treatment.

**Conclusion:** The results suggest that strengthening the immune system during EMT as well as its specialization in the detection of DTCs (disseminated tumor cells) can be effective strategies in the treatment of metastasis.

## INTRODUCTION

Cancer is a generic name given to a group of cells that have lost their specialization and control of their growth ^[1]^. According to the WHO ^[2]^, being one of the main causes of death worldwide, the number of cases is expected to increase in the coming years. It is a system that self-organizes in time and space far from thermodynamic equilibrium, showing high adaptability, resistance and plasticity ^[1]^.

Cancer groups more than 200 diseases, which have common characteristics such as: cells with resistance to apoptosis, induction of angiogenesis, sustained signals of cell proliferation, evasion of growth suppressants, replicative immortality, active invasion and metastasis ^[3, 4]^. Metastasis is the major cause of death in the vast majority of cancer patients ^[5]^. However, the mechanisms underlying each step of this complex process remains obscure.

In metastasis, the tumor usually invades the surrounding tissue and distant points in the body ^[6,7]^. Metastasis presents key stages in which the body manages to fight the disease. These points are presented as targets to be used as treatments ^[8]^. Among these stages are: Epithelial-Mesenchymal Transition (EMT), extravasation and colonization among others ^[8–10]^.

EMT is a process that evades the immune system ^[11]^, to perform invasion and colonization of adjacent and distant tissues ^[12]^. EMT plays a fundamental role in embryonic development and metastasis ^[12]^. During the EMT process, cell-cell and cell-basement membrane interaction is lost. Cells lose the shape and typical polarity of the epithelial phenotype through genotypic changes ^[13]^. These cells acquire a mesenchymal phenotype that is characterized by high invasive capacity and resistance to apoptosis ^[14]^. EMT has the ability to modify cells of the immune system in the tumor microenvironment ^[15]^.

In previous works we have shown that the growth of cancer studied through cell population models and interactions with immune cells, is a complex process. Control in such process could be precisely in that interaction with the immune system; this could be the source of its complexity and its adaptability ^[15,16]^. In this sense, a model was proposed that simulates the effect of combined therapies (chrono-immunotherapy) for the treatment of cancer in the metastatic stage ^[17]^.

The goal of this work is to propose two empirical models that rescue the essential characteristics of EMT and its link with metastasis, as well as a series of key stages, which are not favored in metastasis, which leads to delineating future ways to improve efficacy of treatments aimed at the metastasis process.

## METHODS

Mathematical models represent a suitable procedure for formalizing the knowledge of living systems obtained through theoretical biology ^[18,19]^. Mathematical modeling of tumor metastasis makes possible the description of regularities and it is useful in providing effective guidelines for cancer therapy, drug development, and clinical decision-making ^[19, 20]^.

To carry out the mathematical modeling, the classical method of chemical kinetics was applied, which applies the law of mass action to a mechanism and to obtain a system of ordinary differential equations (ODEs).

Fixed points, stability analysis and bifurcations were calculated using the standard procedure ^[21–24]^. Lyapunov exponents were calculated using the Wolf algorithm ^[22]^. Lyapunov dimension *DL*, also known as Kaplan–Yorke dimension ^[25]^, was evaluated across the spectrum of Lyapunov exponents *λ*_*j*_ as:

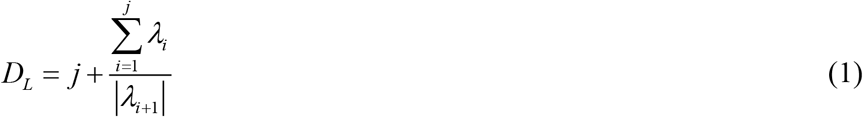

where *λ*_*j*_ is the largest integer number for which *λ*_*1*_ + *λ*_*2*_ + … + *λ*_*j*_ ≥ 0.

COPASI v. 4.6.32 ^[26]^ software was used. However numerical integration was performed on the system of ODEs.

## RESULTS

Tumor cells with epithelial phenotype composing the tumor receive the signal of duplication through growth factors ^[27]^. These cells begin to make the phenotype change due to external factors, such as the effect of the microenvironment ^[28]^ and/or interaction with the immune system ^[29]^. In 2017 Takigawa *et al.* ^[30]^ stated that mesenchymal stem cells induce epithelial to mesenchymal transition in colon cancer cells through direct cell-to-cell contact.

While transitioning between the epithelial and mesenchymal phenotypes, cells can stabilize hybrid epithelial/mesenchymal (E/M) phenotype. Cells in this phenotype have mixed epithelial and mesenchymal properties, which allow them better adherence than mesenchymal cells and better migration than epithelial cells, thereby allowing them to move collectively as clusters. If these clusters reach the bloodstream intact, they can give rise to clusters of circulating tumor cells (CTCs), as have often been seen experimentally ^[31]^.

Based on the previous statements, we propose a model that takes into account EMT, according to the network structure shown in Fig. 1.

**Figure 1.**
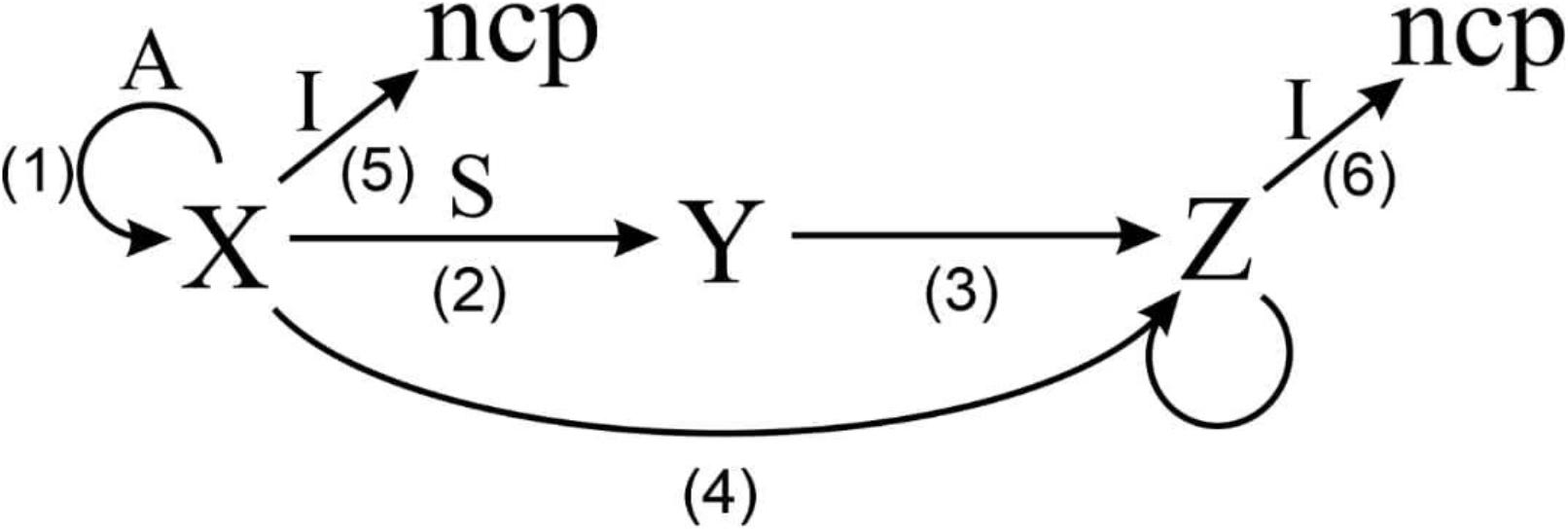
Model network of EMT. *A* represents the action of growth factors in the duplication process; *S* represents the action of external factors al at the beginning of metastasis; *I* represent the population of immune cells; *x* represent the population of epithelial tumor cells, *y* represent the population of hybrids tumor cells and *z* represent the population of mesenchymal tumor cells.

In the model *A* represents the action of growth factors in the duplication process and *S* represents the action of external factors al at the beginning of metastasis ^[32]^; *I* is the population of immune cells (T lymphocytes (CTL) and natural killer (NK)) ^[30]^, *A* and *S* are considered as constants due to the fact that they quantify effects of internal and external factors; we posit *I* as the control parameter ^[34]^ because the population of immune cells may increase or decrease. Variables: *x*, *y*, *z* represent the population of epithelial, hybrids and mesenchymal tumor cells, respectively. Finally, *ncp* represents a non-cancerous product due to the action of immune cells.

Step 1 is related to the mitosis process aided by growth factors; step 2 is related to the beginning of the MET until the formation of the hybrid phenotype, assisted by external transition factors such as snail ^[29]^, step 3 is the completion of the transition, from the hybrid phenotype to the mesenchymal phenotype; step 4 is related to induction of EMT to epithelial cells by mesenchymal stem cells ^[21].^ Finally steps 5 and 6 show the action of immune cells *I*.

The constant values related to each step (see Fig. 1) were chosen empirically trying to achieve the greatest generality and simplicity possible, so we have: *k*_1_ = 4.7 ^ml^/_mmol.s_, *k*_2_ = 1 ^ml^/_mmol.s_, *k*_3_ = 1 s^−1^, *k*_4_ = 1 ^ml^/_mmol.s_, *k*_5_ = 2 ^ml^/_mmol.s_ and *k*_6_ = 2 ^ml^/_mmol.s_.

After these considerations, the system of ordinary differential equations (ODEs) which describes the first steps of the cancer metastasis, according to the proposed model (Fig. 1), has the form:

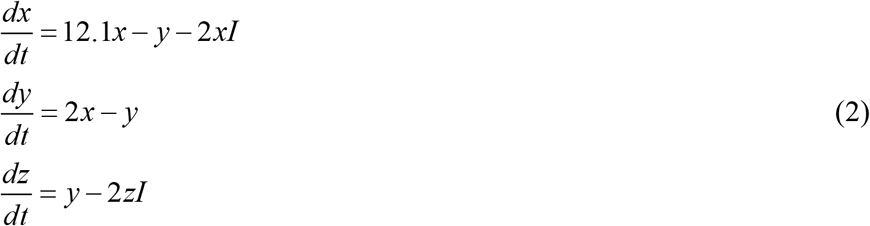

Table 1 shows the numerical results obtained from the stability analysis and the Lyapunov exponents, for two of the values of the control parameter.

**Table 1.** Stability, and complexity for the system of ODEs for different values of the control parameter *I* (*A* = 3, *S* = 2).

| $I$ | Eigenvalues of the Jacobian matrix | Lyapunov exponents $\lambda_i$ |
| --- | --- | --- |
| 3 | -0.05 -2.3i | -0.005 |
| $ss_s$ | -0.05 +2.3i | -0.005 |
| Stable focus | -6.899 | -6.896 |
| 1 | 0.24 -2.4i | $\sim 0.0$ |
| Limit cycle | 0.24 +2.4i | -0.923 |
| (saddle-foci) | -5.078 | -7.054 |

Fig. 2 shows the dynamical behavior of the proposed chemical network model for different values of the control parameter *I* (from 0 to 3). At *I* = 3, there is a stationary state. As *I* value decreases (Fig. 2), the system presents the same dynamic behavior but different values of fixed points. At the critical point *I* = 2.3, dynamics change to a limit cycle. Thus, the dynamical behavior turns oscillatory (Table 1).

**Figure 2.**
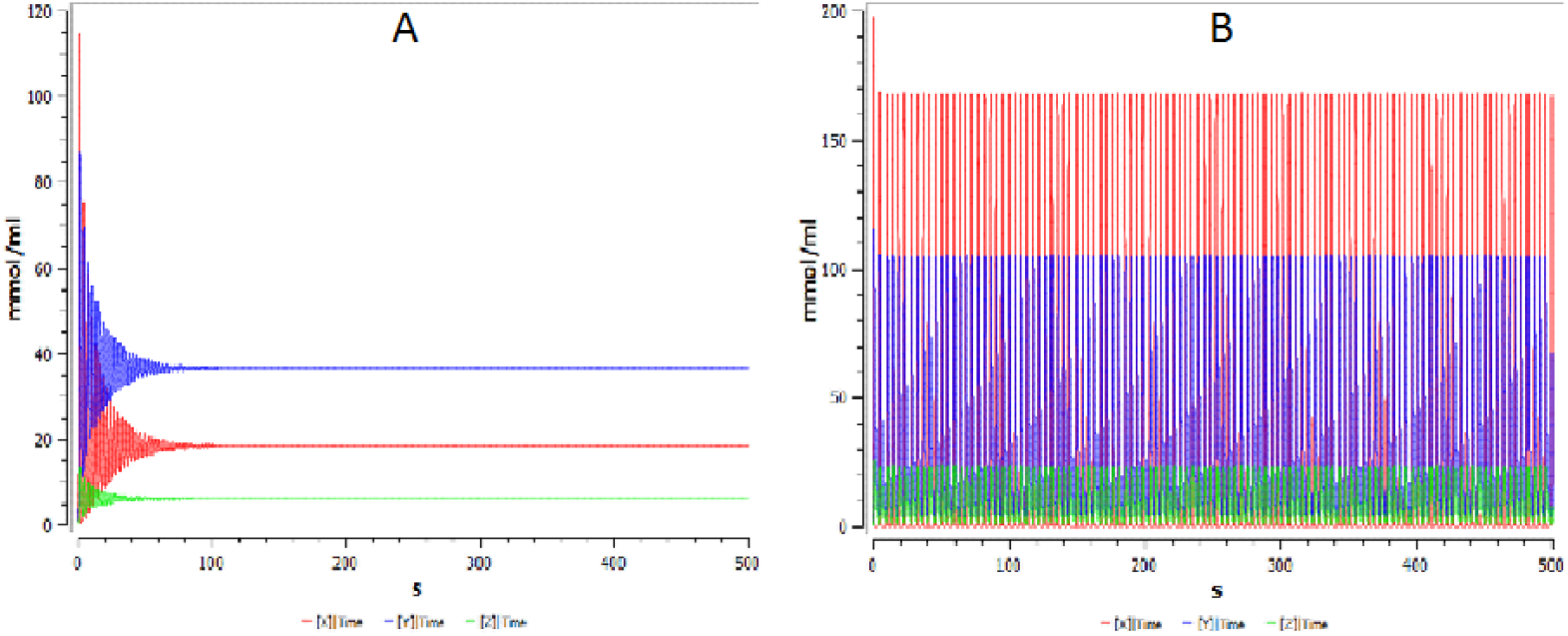
Dynamical behavior of the proposed model for different values of the control parameter *I* (*A* = 3, *S* = 2) : (A) stationary state (*I* = 3), (B) limit cycle (*I* = 1.8); red (population of epithelial tumor cells *x*), blue (population of hybrids tumor cells *y*) and green (population of mesenchymal tumor cells *z*).

When cancer is in the stage of metastasis, it uses the EMT in order to circumvent the body's immune surveillance, as one of its strategies for the proliferation of the primary tumor and a secondary one ^[21]^.

Extremely inefficient processes occur in the final stages of metastasis, such as extravasation and colonization, for example while numerous circulating tumor cells (CTCs) are detected in the blood, disproportionally few metastatic cells are clinically detectable ^[18]^

It is known that in these later stages of metastasis, cells have to face a new hostile microenvironment to reach a pre-metastatic niche ^[34]^. Niche formation was originally described as an accumulation of myeloid cells expressing Vascular Endothelial Growth Factors (VEGFs) at distant metastatic sites before tumor cells arrive ^[35]^ and it is characterized by the recruitment of neutrophils and macrophages ^[36]^.

Most tumors release millions of cells into the bloodstream, but only a small number of metastatic lesions develop, indicating the inefficiency of the metastasis process ^[37]^, this means weakness of metastasis, despite the fact that, as has been shown in previous work ^[16]^, metastasis is a highly robust process. In the extravasation process, tumor cells undergo changes to improve adhesion; this process is closely related to the populations of the immune system and to the action of the new microenvironment ^[37]^. Once they reach the niche, these cells can remain for long periods of time in a dormant state, which makes these populations resist chemotherapies ^[38, 39]^.

It is unknown how the mechanisms of escape from tumor dormancy influence the survival and development of secondary tumors ^[40]^. Recent work proposes evidence of a new inflammatory mechanism produced by immune cells that causes tumor cells in the lung to depart from dormancy to a more aggressive metastasis ^[39]^.

Despite this, the process of metastasis, as stated before ^[5,6]^, constitutes the crucial stage in the evolution of cancer and is responsible for the causes of death. In this sense, a second model is proposed that collects the characteristics of cell populations in these stages of metastasis, such that it reflects its weaknesses, which can be used as potential targets in treatment ^[42]^.

The following empirical model (see Fig.3) compile in a general way, the most important aspects of the extravasation and invasion processes. In this model *P* represents the population of disseminated tumor cells (DTCs) in metastatic niches, *L* represents the population of tumor cells in a dormant state, *R* represents the population of replicative cells once they leave the dormant state and *I* represents the population of cells of the immune system (NK cell, T lymphocytes, neutrophils and macrophages, etc.).

**Figure 3.**
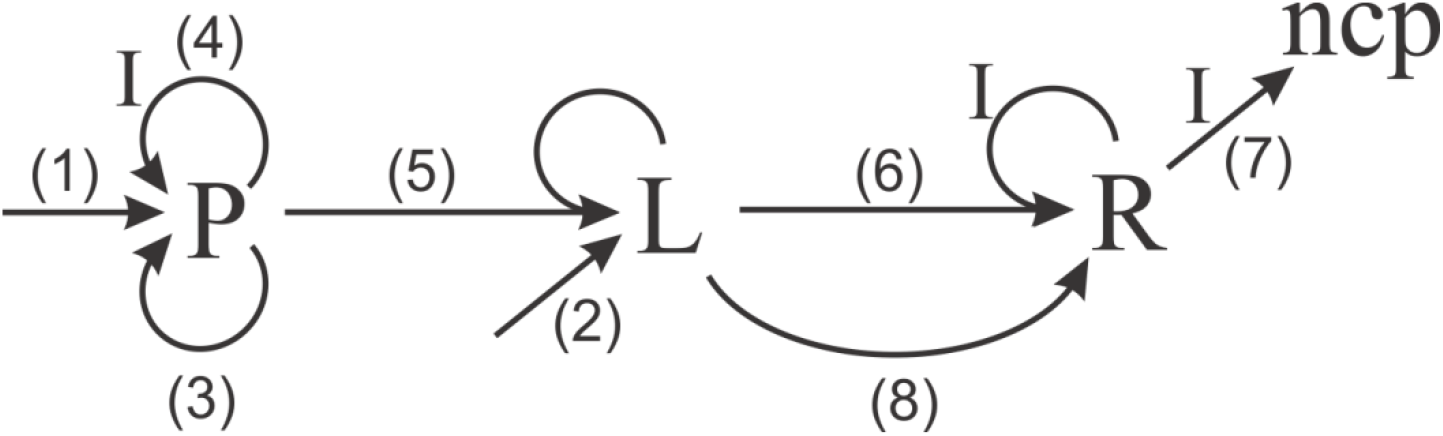
Model network of latest steps of metastasis. *P* represents the population of disseminated tumor cells (DTCs) in metastatic niches, *L* represents the population of latent disseminated tumor cells, *R* represents the population of replicative cells once they leave dormancy and *I* represents the population of cells of the immune system.

*I* value is taken as constant (*I* = 1). Finally, *ncp* represents a non-cancerous product due to the action of immune cells.

Steps 1 and 2 are related with the process of arrival of cancer cells to the niche, Depending on the conditions of this niche, this population may proliferate or enter a dormant state. Steps 3 and 4 are related to the beginning of proliferation and the action of the immune system to inhibit duplication (NK cell, T lymphocytes) ^[42, 43]^. Step 5 represents the arrival of proliferating cells to the niche where the latent tumor remains. Step 6 is related to the action of the populations of the immune system (neutrophils and macrophages) in leaving from the dormancy state. Step 7 is the action of the immune system (NK cell, T lymphocytes) on the population of replicative cells that came out of the dormancy state. Finally step 8 represents the replication of the population of cells that emerged from the dormancy state.

The constants for the model proposed (see Fig. 3) were chosen empirically ^[6]^ trying to have a greater generality and simplicity as possible, so we have: *k*_1_ = 0.1 ^ml^/_mmol.s_, *k*_2_ = 0.1 ^ml^/_mmol.s_, *k*_3_ = 1 s^−1^, *k*_4_ = (0.45 − 0.1) ^ml^/_mmol.s_, *k*_5_ = 1 ^ml^/_mmol.s_, and *k*_6_ = 1 ^ml^/_mmol.s_, *k*_7_ = 0.001 ^ml^/_mmol.s_, *k*_8_ = 24 ^ml^/_mmol.s_, *k*_9_ = 10 ^ml^/_mmol.s_.

According to the proposed model (Fig. 3), the system of ordinary differential equations (ODEs) which describes the extravasation and colonization steps of the cancer metastasis, has the form:

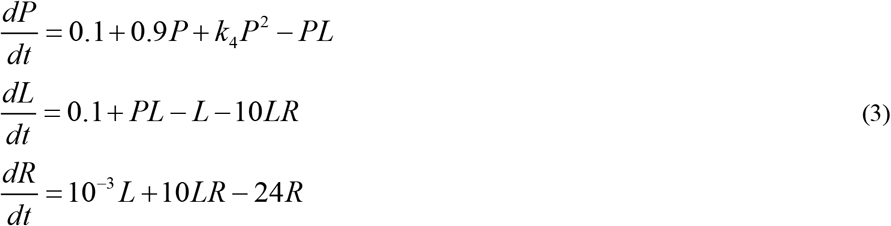

DTCs in the secondary site microenvironment are known to evade immunevigilance of T lymphocytes and NK cell ^[42, 43]^. Furthermore, in EMT, the system self-organizes for a low population of the immune system, as shown in the previous model (see Fig. 1). That is why it is used a constant value for *I* (*I* = 1) and *k*_4_ is chosen as a control parameter since it reflects the dynamics of the immune system in the growth of the secondary tumor. Thus, as the constant *k*_4_ increases its value, the action of the immune system in tumor development decreases.

Table 2 shows the numerical results obtained from the stability analysis, the Lyapunov exponents and the Kaplan-York dimension, for different values of the control parameter.

**Table 2.**
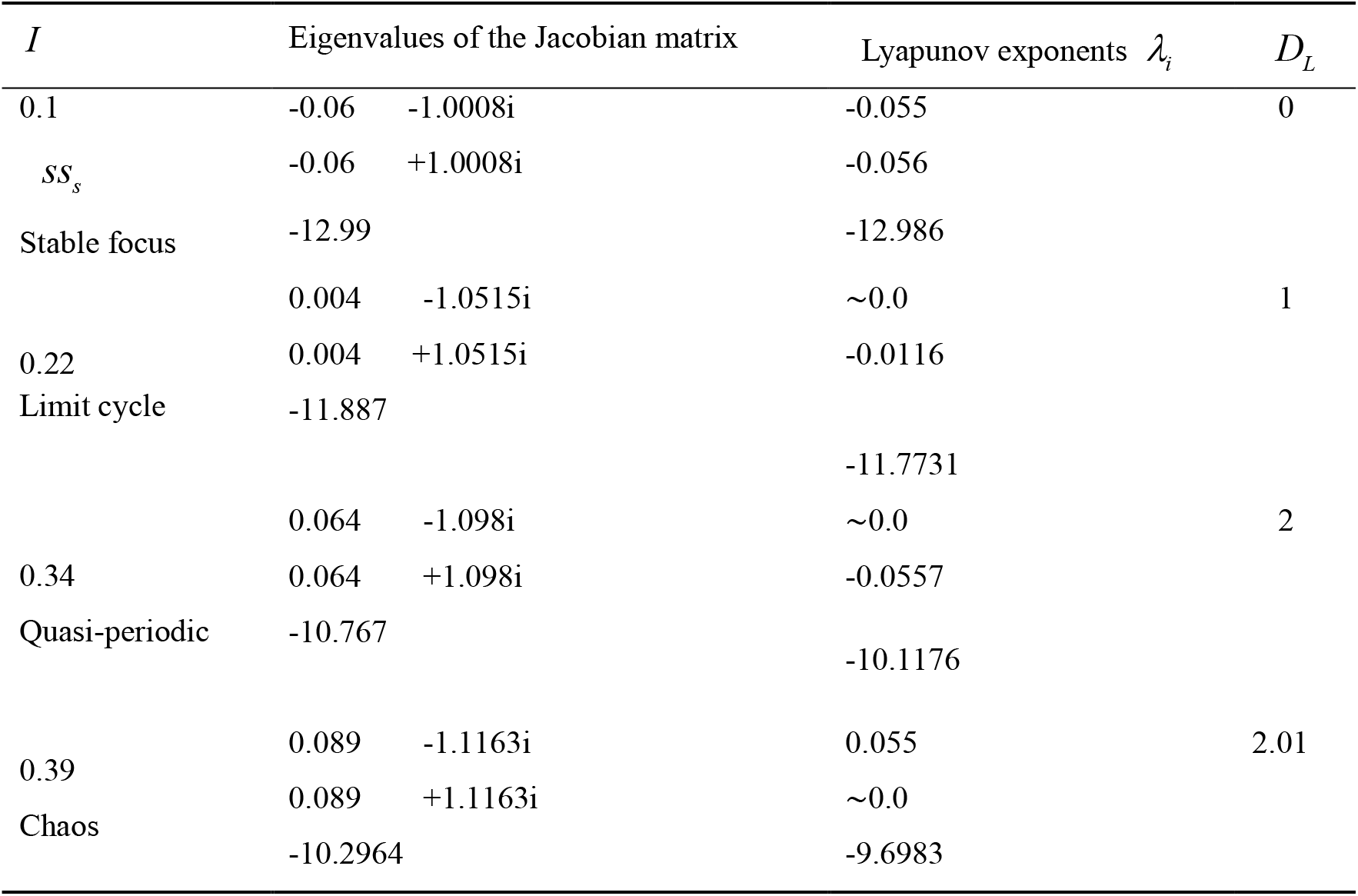
Stability, and complexity for the system of ODEs for different values of the control parameter *k*_4_ (*I* = 1).

Fig. 4 show the dynamical behavior of the proposed network model for different values of the control parameter *k*_4_ (from 0.45 to 0.1). At *k*_4_ = 0.1, there is a stationary state. As *k*_4_ is increasing the system presents the same dynamic behavior but different values of fixed points. However, at the critical point *k*_4_ = 0.22, dynamic behavior changes to a limit cycle. Thus, it turns oscillatory. As *k*_4_ continue decreasing and reaches *k*_4_ ≈ 0.34, a new qualitative change occurs: the limit cycle suffers a distortion: there exists two maxima for the values of each the oscillating variables (*P*, *L*, *R*). And so on, a cascade of bifurcations is triggered. The chaotic dynamics is reached at *k*_4_ ≈ 0.39 (see fig. 4 D, Table 2).

**Figure 4.**
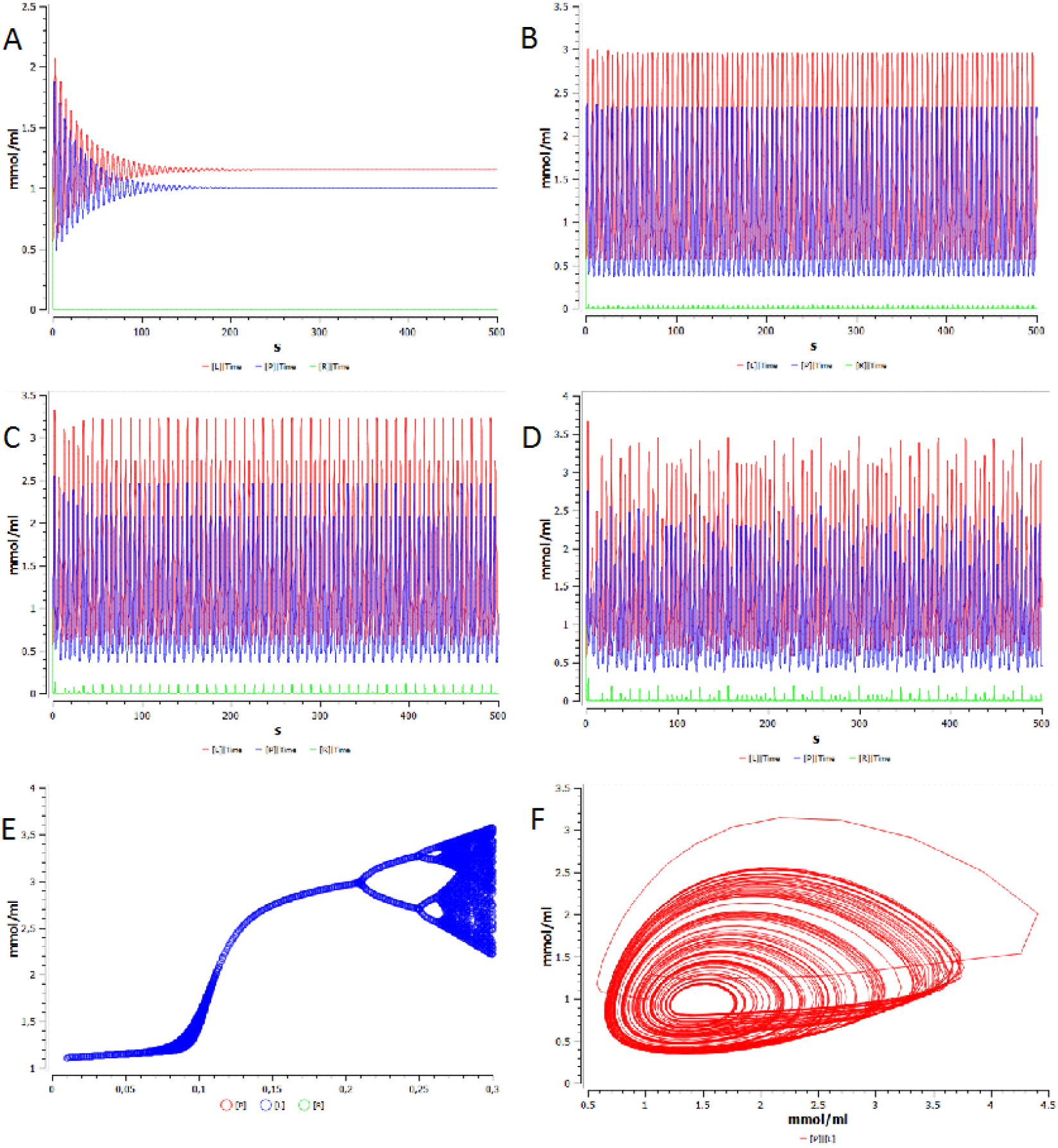
Dynamical behavior and bifurcation diagram of the proposed model for different values of the control parameter *k*_4_ (*I* = 1): (A) stationary state (*k*_4_ = 0.1), (B) limit cycle (*k*_4_ = 0.15), (C) quasiperiodic (*k*_4_ = 0.23), (D) Chaos (*k*_4_ = 0.35), (E) route to chaos, (F)-chaos like attractor; Red corresponds to L populations (cells in dormant state), blue color corresponds to P population (DTC and proliferative cells) and green corresponds to R (cells out of dormant state).

## DISCUSSION

As previously mentioned, one of the processes by which cancer in the metastatic stage evades the immune system is EMT. The system self-organizes outside the thermodynamic equilibrium in an oscillatory regime (fig. 2 B). It confers greater robustness and therefore adaptability and resistance to therapies ^[44, 45]^. In figure 2 it is observed that, as the population of cells of the immune system increases, the process goes to a stable steady state (fig. 2 A). On the other hand it is observed that, when the system loses self-organization, the population of hybrids tumor cells decreases as the systems display a hybrid epithelial-mesenchymal configuration, thus what is reported in the results suggests that EMT is rarely an all-or-nothing process ^[46]^. Tian *et al.* ^[47]^ stated how increasing the T cell population leads to decreased EMT. It is therefore that a combined immune therapy and directed to the EMT, would lead to decrease the robustness of the system and consequently to increase the efficacy in the treatment of tumors in advanced phase. Resistance to therapeutic regimens is a key problem for cancer therapy, as it precludes complete remove of the tumour and enables tumour recurrence, which is one of the main causes of death ^[10, 41]^.

The colonization and outgrowth of tumor cells in a secondary organ is often considered the rate-limiting, as well as the most poorly delineated step in the metastatic cascade ^[29]^. Pre-metastatic niche theory shows that before arriving disseminated tumor cells (DTC), bone marrow-derived haematopoietic stem cells are recruited by tumour-derived factors to the secondary site where they transfer a more hospitable microenvironment to foster the survival and expansion of metastatic lesions ^[40]^. According to the self-seeding hypothesis, metastatic tumor cells can also return to the primary site, accelerating the growth and malignant evolution of primary tumors ^[48]^.

Metastasis is known to be a highly complex process (see Figure 4D), which overlaps the immune system ^[27]^. In this model, we observe how the System behaves when varying the growth rate of DTCs. The appearance of chaotic dynamical the model could explain the relapses and poor prognosis of the disease. Such behavior maybe gives a higher degree of robustness, and the possibility of creating new information and learning ability.

This is why the design of a therapy that focuses, at this stage, on enhancing the immune action focused on the growth of DTCs in the niche will have greater efficacy. As seen in Figure 4, as the control parameter decreases, the population of disseminated tumor cells will decrease, leading to a decrease in the complexity of the metastatic lesion. In other words, this process is related to the increased action of the immune system. This would lead the system to lose self-organization (see fig. 4A), leading to a less robust steady state (See Table 2).

At this stage of metastasis, dormant tumors (L) in secondary sites also play an important role in tumor regression after surgical recession ^[49]^. In addition, cells when leaving the latency (R) show a more aggressive behavior ^[50]^. This is why self-organization at this stage makes resistance to therapy more pronounced. ^[17]^ In other words, becomes more robust and consequently exhibits a greater adaptation to the conditions of the environment.

In this work, we propose models that represent tumor metastasis: a process out of thermodynamic equilibrium that goes through different stages. From the stages contained in the previous discussion, we postulate that EMT, extravasation, and colonization play a crucial role in the metastasis process. For this, we elaborate two models that reproduce some of the main characteristics.

In summary, in this paper we have found that:

1. A therapy aimed at boosting the immune system in a combined way and directing it to EMT, would lead to decrease the self-organization, by decreasing complexity and consequently robustness of the system. This should lead to increase the efficacy in the treatment of tumors in advanced phase. This process gives the system a route to evade the immune system and it gives the cells in the bloodstream a greater survival potential.
2. 2. A therapy aimed at specializing the immune system against DTCs could reduce resistance and adaptability of tumors in the last stages of the metastatic cascade, making the system less robust.

The current theoretical framework will hopefully provide a better understanding of cancer and contribute to improvements in cancer treatment.

## DECLARATIONS

## Acknowledgments

Prof. Dr. A. Alzola and Prof. Dr. Germinal Cocho *in memoriam*. JMNV thanked the CEIICH of the UNAM Mexico for the warm hospitality.

## Authors’ contributions

All authors contributed equally to the completion of this article.

## Availability of data and materials

Not applicable.

## Financial support and sponsorship

The financial support by PREI-DGAPA-2019

## Conflicts of interest

All authors declared that there are no conflicts of interest.

## Ethical approval and consent to participate

“Not applicable.”

## Consent for publication

Not applicable.

## Copyright

© The Author(s)

## Notes

### Competing Interest Statement

The authors have declared no competing interest.

